# Deep rhizospheres extend the nitrogen cycle meters below the base of soil into weathered bedrock

**DOI:** 10.1101/2024.01.08.574278

**Authors:** Kelsey Crutchfield-Peters, Daniella M. Rempe, Alison K. Tune, Todd E. Dawson

**Affiliations:** University of California, Berkeley, Department of Integrative Biology; University of Texas at Austin, Jackson School of Geosciences

## Abstract

Nitrogen is the most limiting nutrient to forest productivity worldwide. Recently, it has been established that diverse ecosystems source a substantial fraction of their water from weathered bedrock, leading to questions about whether root-driven nitrogen cycling extends into weathered bedrock as well. In this study, we specifically examined nitrogen dynamics using specialized instrumentation distributed across a 16 m weathered bedrock vadose zone (WBVZ) underlying an old growth forest in northern California where the rhizosphere—composed of plant roots and their associated microbiome—extends meters into rock. We documented total dissolved nitrogen (TDN), dissolved organic carbon (DOC), inorganic N (ammonium and nitrate) and CO_2_ and O_2_ gasses every 1.5 m to 16 m depth for two years. We found that biologically available nitrogen in the weathered bedrock rhizosphere was comparable in concentration to temperate forest soils and primarily organic. TDN concentrations in the WBVZ exhibited distinct patterns with depth and were correlated with periods of increased whole-ecosystem metabolic activity as well as stream discharge, suggesting competing rhizosphere and leaching processes in the fate of TDN in the WBVZ. Carbon isotope composition of the DOC suggests that dissolved organic matter in the WBVZ is primarily derived from fresh plant sources. We conclude that N cycling in the WBVZ is driven by an active rhizosphere meters below the base of soil and represents an important and overlooked component of deeply rooted ecosystems that must be incorporated into future models and theory of ecosystem function.

## Introduction

Plant roots and their associated microbiome–collectively termed the *rhizosphere*–perform almost all plant water and nutrient uptake necessary to sustain carbon (C) fixation. Historically, studies on rhizosphere resource acquisition have focused almost exclusively on the shallowest soil horizons (1), despite evidence of roots extending meters into the subsurface and often into fractured rock (2–5). However, a growing number of studies have documented significant water uptake by plants from weathered bedrock vadose zones (WBVZ)—the unsaturated portion of the subsurface beneath soil and above the groundwater table (6–8)—highlighting the importance of weathered bedrock rhizospheres in plant resource acquisition. The extent to which rhizospheres cycle and acquire essential nutrients in the WBVZ, however, remains unknown (9). Given the widespread evidence for roots in weathered bedrock, it is crucial we understand the role of the WBVZ in the nutrient ecology of deeply rooted systems, especially as deeper resources are likely to be more important in the face of future climate change (10).

Rhizospheres are one of the most biogeochemically active regions of the subsurface, affecting water withdrawal, C inputs (i.e., rhizodeposition) and nutrient cycling that can impact the chemistry of entire catchments (11, 12). One of the key elements cycled and acquired by rhizospheres is nitrogen (N)—a fundamental building block of proteins, nucleic acids, and other biomolecules. Due to the relatively low abundance of biologically available forms of N in natural ecosystems, N is often the most limiting nutrient to primary productivity in terrestrial ecosystems (13). As such, plants have evolved diverse uptake mechanisms to acquire both inorganic (ammonium, NH_4_^+^ and nitrate, NO_3_^-^) and organic N under variable soil conditions (14–16).

N has one of the most complex biogeochemical cycles, and its chemical speciation and bioavailability are shaped by biotic and abiotic processes within the rhizosphere, such as plant and microbial immobilization (17, 18), microbial transformation (19, 20), mineral weathering and ambient chemical conditions (21). These processes interact to determine the fate of N in ecosystems, leading to its accumulation in biomass, short-term storage in dissolved, particulate or litter pools, release from sediments and potential loss through leaching and dissimilatory processes (22). In turn, each of these processes is limited by C and water availability (23). Thus, understanding how the sources and fates of N in the weathered bedrock vadose zone change, as water and subsurface C dynamics do, is key to understanding the future C balance and health of deeply rooted ecosystems.

Several studies have pioneered research on deep N cycling: highlighting the importance of lithologically-derived N to diverse ecosystems (24–26), demonstrating rhizosphere-driven bedrock weathering and N availability in shallow rock fractures (27), and providing the first look into N dynamics meters into fractured rock (28). Collectively, this body of work provides strong evidence for N cycling within the WBVZ and the importance of deep N cycling. But to our knowledge, no study has attempted to describe, quantify, and relate long-term N cycling in a WBVZ to carbon and water cycling where a deep rhizosphere exists. In the present study, we investigate a WBVZ where the rhizosphere extends meters into weathered bedrock, asking: (1) how might rhizosphere processes alter the concentration of available N year-round; and (2) what are the sources and fates of N within the WBVZ?

To address these aims, we leverage a unique Vadose Zone Monitoring System at the Eel River Critical Zone Observatory (ERCZO), the only one of its kind located in an old-growth temperate forest, allowing for sampling of water and gasses every 1.5 m down to 16 m below the ground surface. More than a decade of research at the ERCZO has established that rhizosphere-driven C and water cycling extends meters into weathered bedrock (7, 29) and that water limitations are exacerbated by C fixation being out of phase with water delivery at the site (30). Due to the extensive WBVZ, associated rhizosphere, and previous observations of deep plant-related activity at the ERCZO, we hypothesize that the weathered bedrock rhizosphere may receive and access N via several sources (Fig. 1A: S1–S3). These sources include: (S1) downward leaching of inorganic and organic dissolved N from the overlying soil horizons into fractured rock; (S2) in situ root exudation or organic matter decomposition through microbial activity; and (S3) release of N in the form of NH_4_^+^ from the weathering of N-rich parent material. We hypothesize several potential fates for N as well: (F1) uptake and immobilization of inorganic and organic N in plant and microbial (defined here as bacteria, archaea, and fungi) biomass, (F2) gaseous loss through denitrification, and (F3) loss through leaching downward beyond the rooting zone. We assess these hypotheses by providing the first multi-year record of dynamic N and C cycling at discrete depths throughout a WBVZ home to a deep rhizosphere.

**Figure 1.**
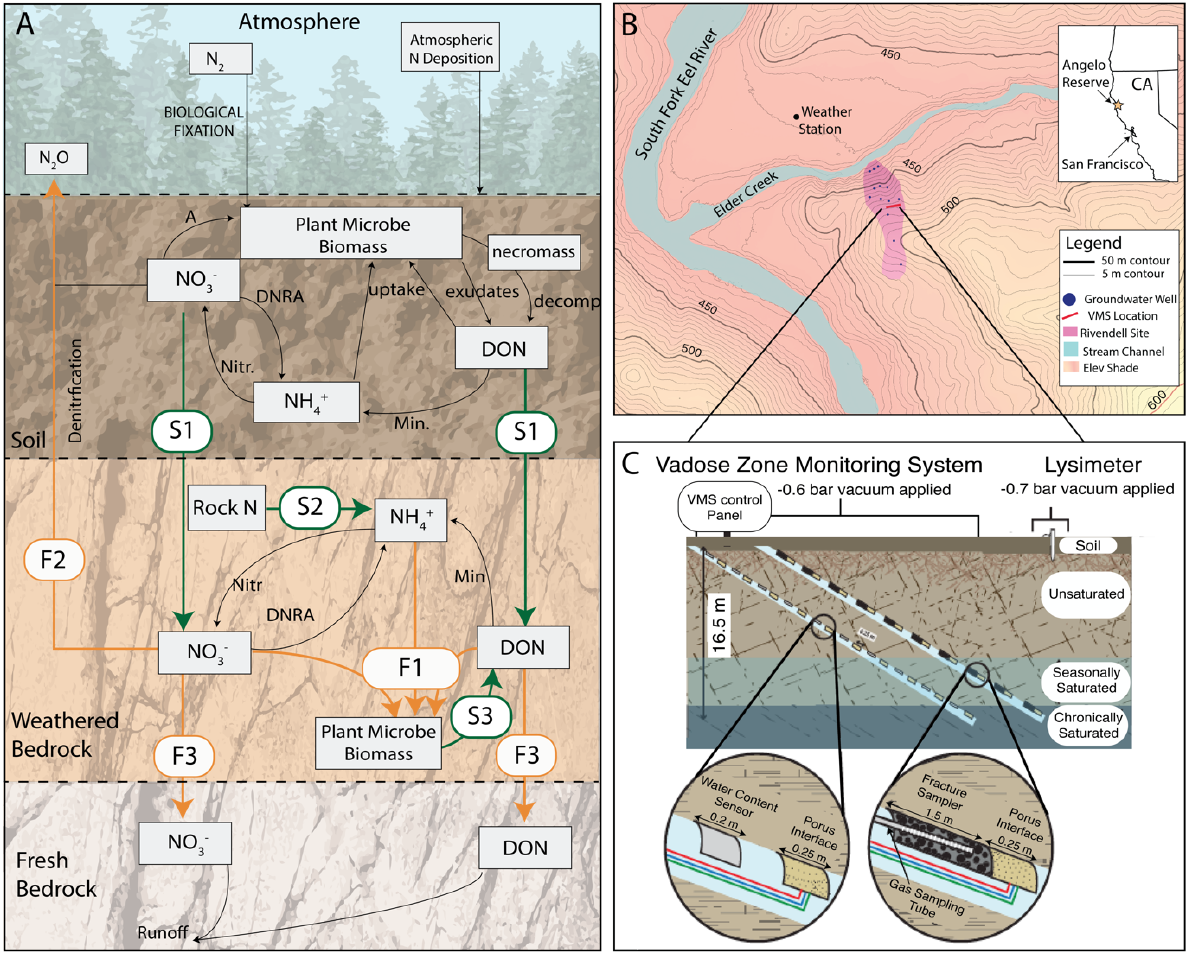
Conceptual diagram of dominant sources (S1–S3), pools, transformations, and fates (F1–F3) of N in the WBVZ (panel A), site map for Rivendell (panel B) and schematic of the vadose zone monitoring system (panel C, modified from Tune et al. 2020). (**A)** Nitr = nitrification, Min = mineralization, DNRA = dissimilatory nitrate reduction to ammonia. (**B)** red line represents the footprint of the VMS. **C** the VMS consists of two 55° diagonal bore holes, each lined with a flexible instrumented sleeve designed to capture freely draining (Sleeve B) and more tightly held (Sleeve A) waters. The entire VMS profile extends 16.5 m below the soil surface and samplers are spaced approximately every 1.5 meters.

## Results

Previous studies leveraging the VMS and groundwater wells across the site provide evidence for an active rhizosphere that extends to approximately 7 m below the ground surface at the ERCZO, resulting in deep water withdrawal and rhizosphere respiration (7, 29). In Figure 2, a profile illustrates the depths of soil (0–0.8 m), weathered bedrock that remains unsaturated (i.e above the water table) year-round (0.8–6.9 m), weathered bedrock that is seasonally saturated during the wet season (7–14.9 m), and bedrock that remains saturated year-round (> 15 m) at the VMS installation location. All evidence suggests that rhizosphere activity is restricted to the chronically unsaturated WBVZ. We therefore consider 0–7 m as the rooting zone in our following analyses on linked N and C in the WBVZ.

**Figure 2.**
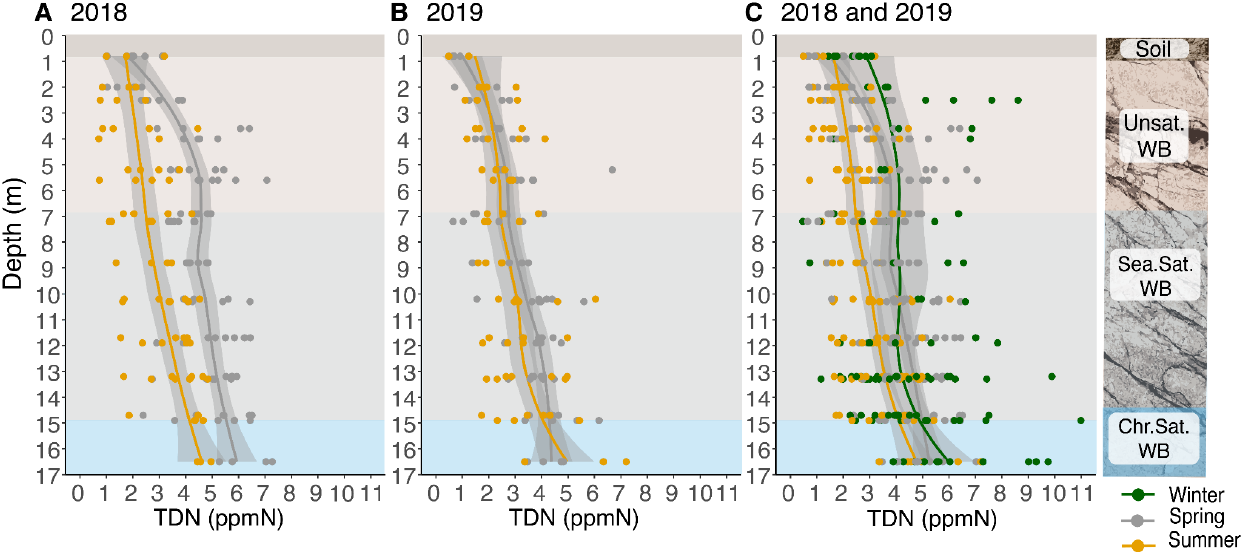
Total dissolved N (ppm, mg/L) in spring (gray) and summer (yellow) months for 2018 (panel A) and 2019 (panel B). Panel C combines 2018 and 2019 and includes TDN from winter months (green) where samples were available. Shaded boxes represent depth horizons that are classified as soil (dark brown, 0-0.8 m), unsaturated bedrock (light brown 0.8-6.7 m), seasonally saturated rock (light gray 6.7-14.2 m) and chronically saturated bedrock (light blue > 14.2 m).

### Ecologically significant quantities of N in the WBVZ

Total Dissolved N (TDN) concentrations within and below the rooting zone were comparable to what is observed in some temperate forest soils (mean = 3.66 ± 1.66 ppm, N = 456, Fig. 2A–C; Fig. S1) (31, 32), and were an order of magnitude higher than what was observed in the limited soil pore water samples from the sampling period (0.28 ± 0.21 ppmN, n = 25). NH_4_^+^ concentrations in the VMS were also comparable to nutrient-poor temperate forest soils (mean = 0.13 ± 0.10 ppm)(32) across depth and season (Fig. 3A), while NO_3_^-^ was frequently below detection year-round (< 0.05 ppm) with periodic pulses in concentration at certain depths (e.g., 5 meters, 10.1 meters and 13.2 meters, Fig. 3B). From TDN and total inorganic N ([NH_4_^+^] + [NO_3_^-^]) measurements, we estimate DON as the dominant form of TDN across all depths year-round, ranging from 77% to 99% (mean = 95.4 ± 4.1%) of all TDN in the WBVZ (Fig. 4A). While it is common for soil TDN to be largely DON, in the only other study of seasonal N dynamics in a WBVZ, inorganic nitrogen was the dominant form (28). Importantly, the aforementioned study was on a system that lacked a deep rhizosphere.

**Figure 3.**
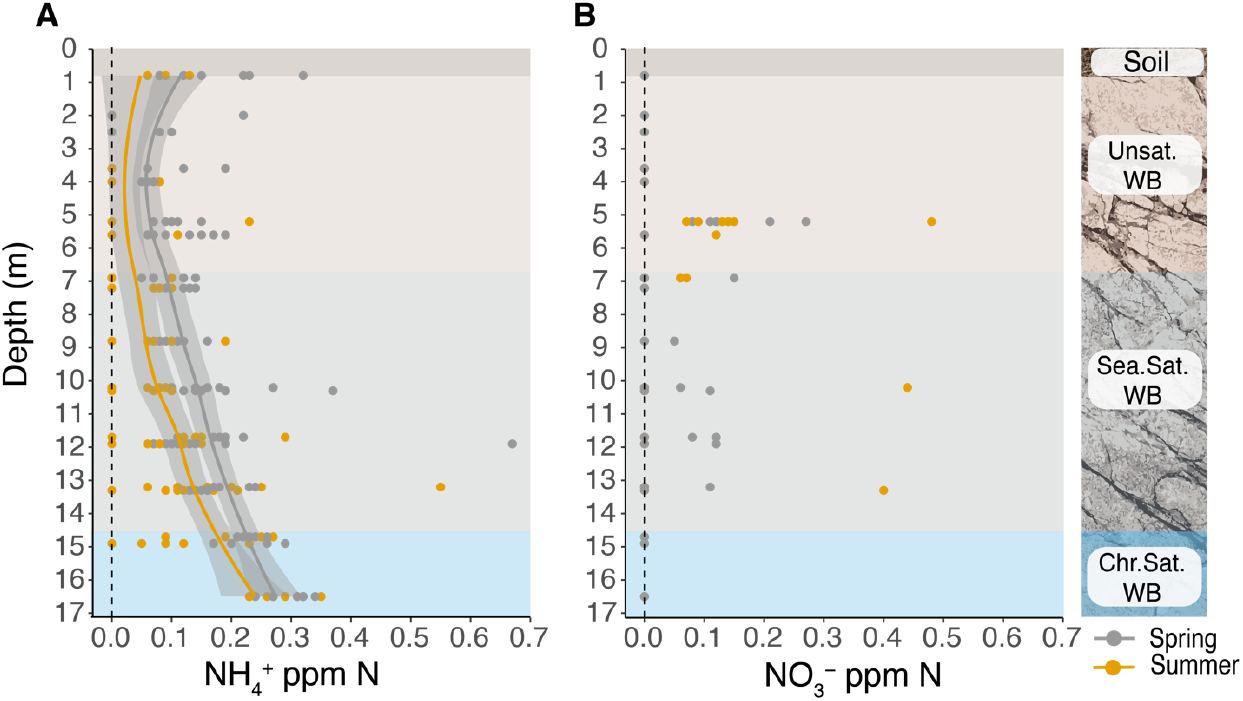
Inorganic N (NH_4_^+^ and NO_3_^-^) profiles for combined 2018 and 2019 spring (gray) and summer (yellow) months. NO_3_^-^ is frequently below detection (< 0.05 ppm). Points along 0 line represent times where NO_3_^-^ was below detection. Shaded boxes represent depth horizons that are classified as soil (dark brown, 0-0.8 m), unsaturated bedrock (light brown 0.8-6.7 m), seasonally saturated rock (light gray 6.7-14.2 m) and chronically saturated bedrock (light blue > 14.2 m).

**Figure 4.**
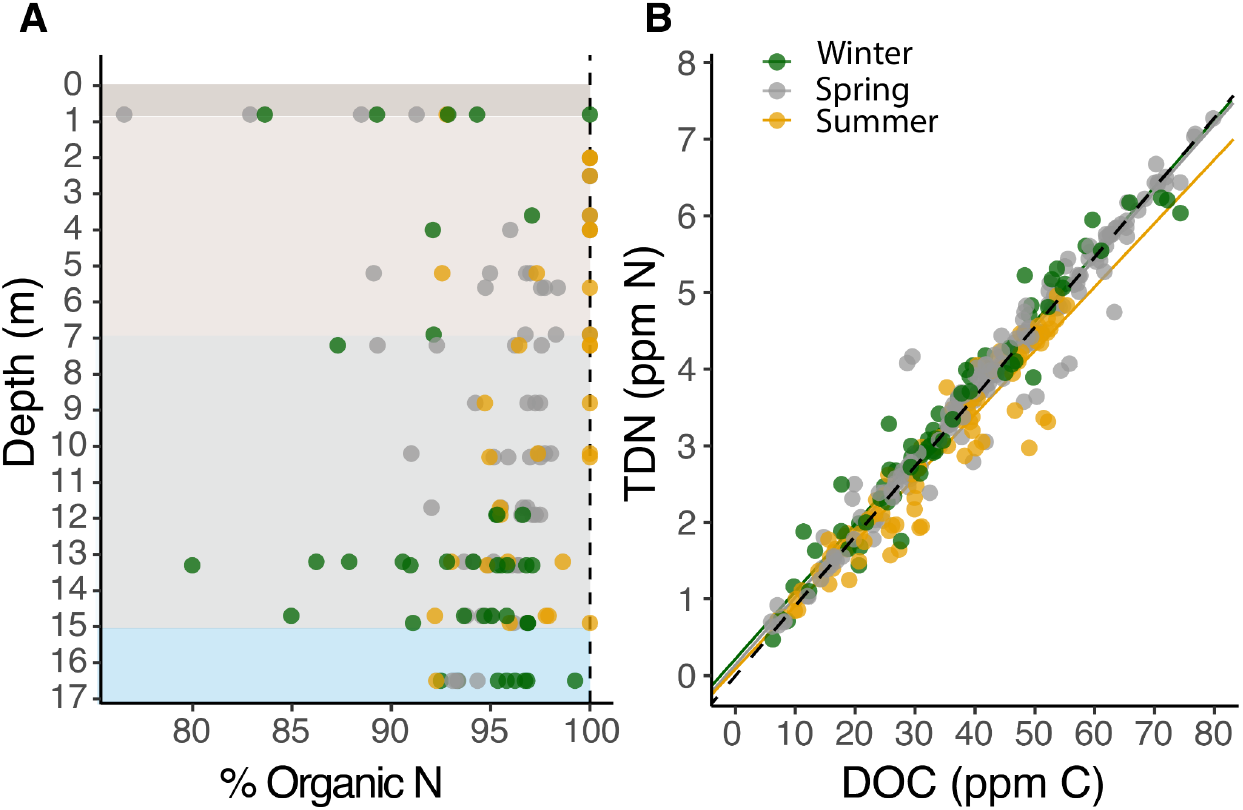
**A** Percent DON (DON = TDN – ([NH_4_^+^] + [NO_3_^-^]). Gray points represent spring (February-April) and yellow points represent summer (May-July). Shaded boxes represent depth horizons that are classified as soil (dark brown, 0-0.8 m), unsaturated bedrock (light brown 0.8-6.7 m), seasonally saturated rock (light gray 6.7-14.2 m) and chronically saturated bedrock (light blue > 14.2 m). **B** and DOC:TDN (panel B) in the VMS. The dashed black line represents a DOC:TDN ratio of 11. Yellow and gray lines represent the summer and spring correlation between DOC and TDN respectively.

### N dynamics respond to depth, season and water year in the WBVZ

Not only did we observe a large quantity of bioavailable N present in the WBVZ, it also exhibited dynamic change in response to various site factors. We used linear mixed models to test the effect of depth, season and water year on TDN. In our full model of TDN concentration in the WBVZ, depth had a strong positive effect on TDN across season and year (p < 0.001; F = 39.7; Table S1), ranging from 1.7 ± 0.87 to 3.30 ± 1.14 ppm N in the rooting zone and increasing to 6.07 ± 1.88 ppm N at the base of the WBVZ (Fig. 2, Table S2). As main effects, both season (p < 0.001; F = 29.9) and water year (p = 0.009; F = 6.8) had a significant effect on TDN, as well as a significant interaction (p < 0.001; F = 28.6).

Post-hoc pairwise comparison of TDN differences by season within-year revealed that TDN was significantly higher in spring 2018 (4.56 ± 0.16 ppm N) than in summer 2018 (2.96 ± 0.18 ppm N), but that there was no significant difference between spring and summer TDN concentrations in 2019 (Fig. 2A–B; Table S2, S3). Winter TDN concentrations exhibited a higher degree of variability in concentration than spring and summer months, as well as a smaller sample size due to naturally low water availability in early winter within the VMS. Nevertheless, TDN reached its highest concentrations in winter 2019 (4.69 ± 0.218 ppm N) and was significantly higher than both spring and summer months. However, TDN in winter of 2018 was significantly lower than spring and only marginally higher than summer concentrations (Table S3). At the site, transition from spring to summer months marks an increase in vegetation transpiration and respiration at the site (29), therefore, decreases in TDN from spring to summer in 2018 may result from increased rhizosphere metabolic activity and uptake within the 7 m rooting zone—reflected in CO_2_ and O_2_ concentrations in the shallow weathered bedrock rooting zone (Fig. S5). However, the absence of a detectable decrease in TDN concentrations between spring and summer 2019 is likely the result of major differences in timing and quantity of precipitation received between 2018 and 2019.

Approximately 920 mm more precipitation was received in 2019 (2570 mm) compared to 2018 (1650 mm), most of which (901 mm) fell during winter months (Fig. S3). To further explore the effect of rainfall on TDN, we examined the relationship of TDN to discharge recorded downstream at the South Fork Eel gauging station as an integrated measure of rainfall received at the site. When accounting for depth effects, we found that discharge exhibited a significant positive effect on TDN (p = 0.013, F = 6.2; Fig. 5), indicating that during times of high precipitation and transport of water to the stream, TDN tends to increase in concentration in the WBVZ. This is apparent in early winter 2019 where TDN spikes in January following significant rains (Fig. 5) and then decreases in following sampling events. This is quite different from 2018 where TDN concentrations gradually increase starting from winter into spring and remain constant until April at which point concentrations gradually decline into June (Fig. 5). Notably, this systematic decline in TDN across depth occurs several weeks after the final rain event of 2018 (April 6) and continues while the water table is declining, suggesting that downward leaching is not driving TDN loss. This further supports plant and microbial uptake as the primary driver of this decrease in TDN within the WBVZ in 2018.

**Figure 5.**
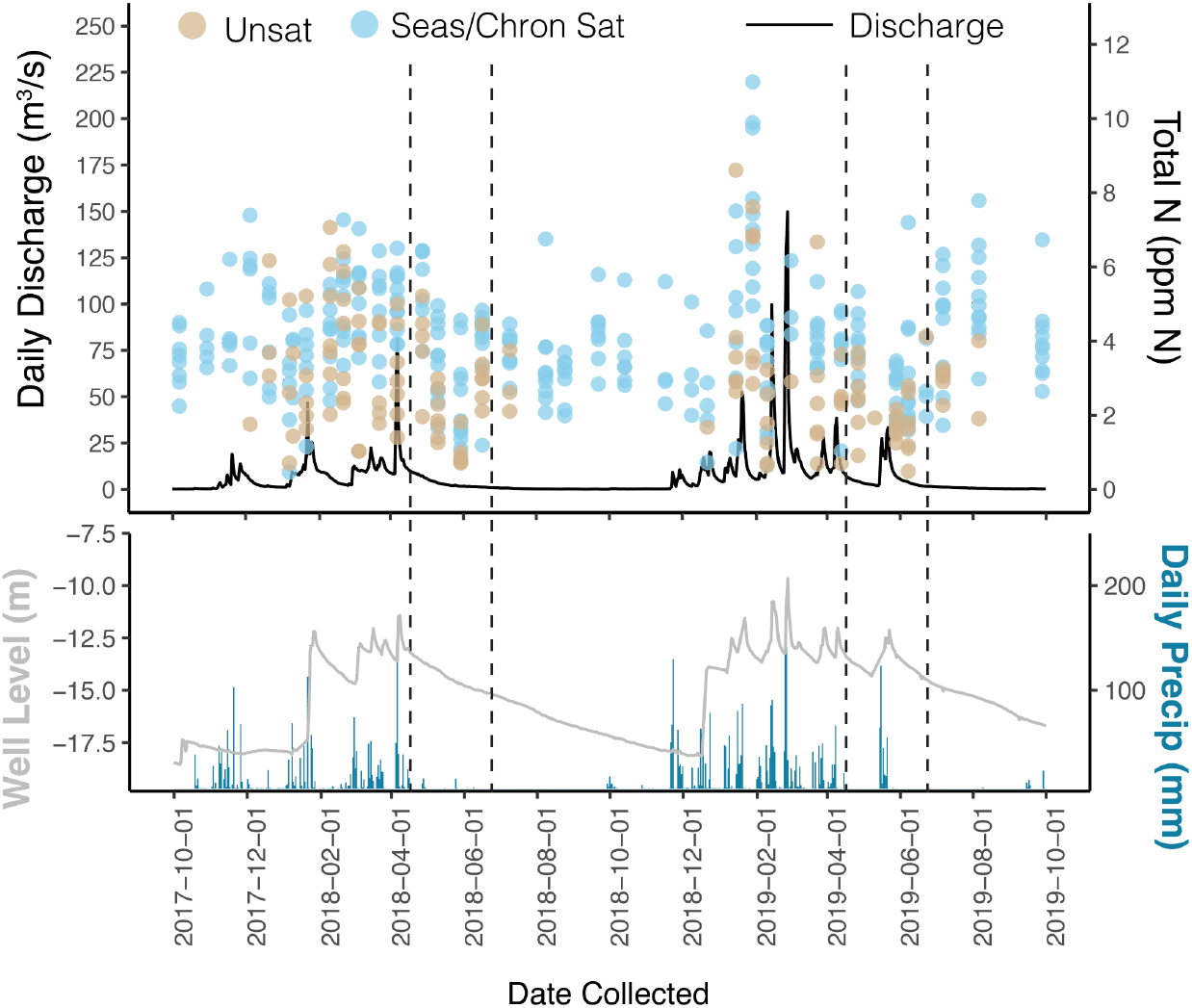
**Top panel:** Total N in the unsaturated (tan points), seasonally saturated and chronically saturated (light blue points) weathered bedrock vadose zone and stream discharge across the sampling period. **Lower panel:** Daily precipitation (blue bars) and water table (well level, m) below the ground surface (gray). Dashed lines represent the bounds between spring and summer where TDN decreased in 2018, but not in 2019. Notably, precipitation continues through this sampling period in 2019 but not in 2018, where the water table continues to decline.

NH_4_^+^ also exhibited temporal and spatial variation in the WBVZ. Like TDN, NH_4_^+^ concentrations tend to increase with depth (p < 0.001 ; F = 35.2; Fig. 3A, Table S4). Unlike TDN, however, the lowest concentrations of NH_4_^+^ are seen at the intermediate depths of the rooting zone, likely indicating uptake by plant roots and their associated microbes. Another difference in dynamics between TDN and NH_4_^+^ occurs in the shallowest portion of the rooting zone, where concentration of NH_4_^+^ spikes upon the first rains, while TDN does not. Season also has a significant effect on NH_4_^+^, with spring months having higher NH_4_^+^ concentrations than summer months, on average (p < 0.001; F=28.4; Fig. 3A). And while NO_3_^-^ is often below detection in the VMS, it does exhibit periodic pulses in concentrations at specific depths (Fig. 3B), suggesting NO_3_^-^ is consistently low in this system and possibly generated and lost rapidly within the WBVZ via leaching. Sampling for inorganic N began in July of 2018, so we were unable to compare inter-annual effects of the 2018 and 2019 water years.

### Plant-derived organic matter is the dominant source of N in the WBVZ

TDN and DOC across all depths and sampling dates exhibit a very strong positive relationship (R^2^ = 0.95, Fig. 4B). This strong relationship results in a consistent DOC:TDN ratio that remains relatively invariant across depth (DOC:TDN = 11.2 ± 1.32; Fig. 2B). The strong correlation between DOC and TDN across depth indicates that the measured C and N are likely linked in the same molecules. Based on this assumption, we use DOC-δ^13^C to trace the source of dissolved organic matter (i.e., linked DOC-DON dynamics) in the WBVZ. To investigate the source of dissolved organic matter in the WBVZ, we compared the stable carbon isotope composition (δ^13^C) of DOC in the VMS to that of bulk solids (including foliage, root tissues, soil, and weathered bedrock; Fig. 6) and soil pore water DOC. We found that δ^13^C-DOC values in VMS waters ranged from -29‰ to -28‰. These values fall within the measured range of δ^13^C for the dominant vegetation at the site (−31‰ to -27‰), and are depleted by several permil relative to soils (−26‰ to -24‰) and bedrock (−25‰ to -24‰). Soil pore water δ^13^C-DOC fell between (−27‰ to -24‰) and tracked quite closely to bulk soil δ^13^C, which became more enriched with depth, as is commonly observed in soil profiles (33). This enriched signal also appears in the shallowest port of the VMS (−24‰ in port B1) that spans the soil and saprolite interface, likely due to leached soil pore water into the shallow WBVZ.

**Figure 6.**
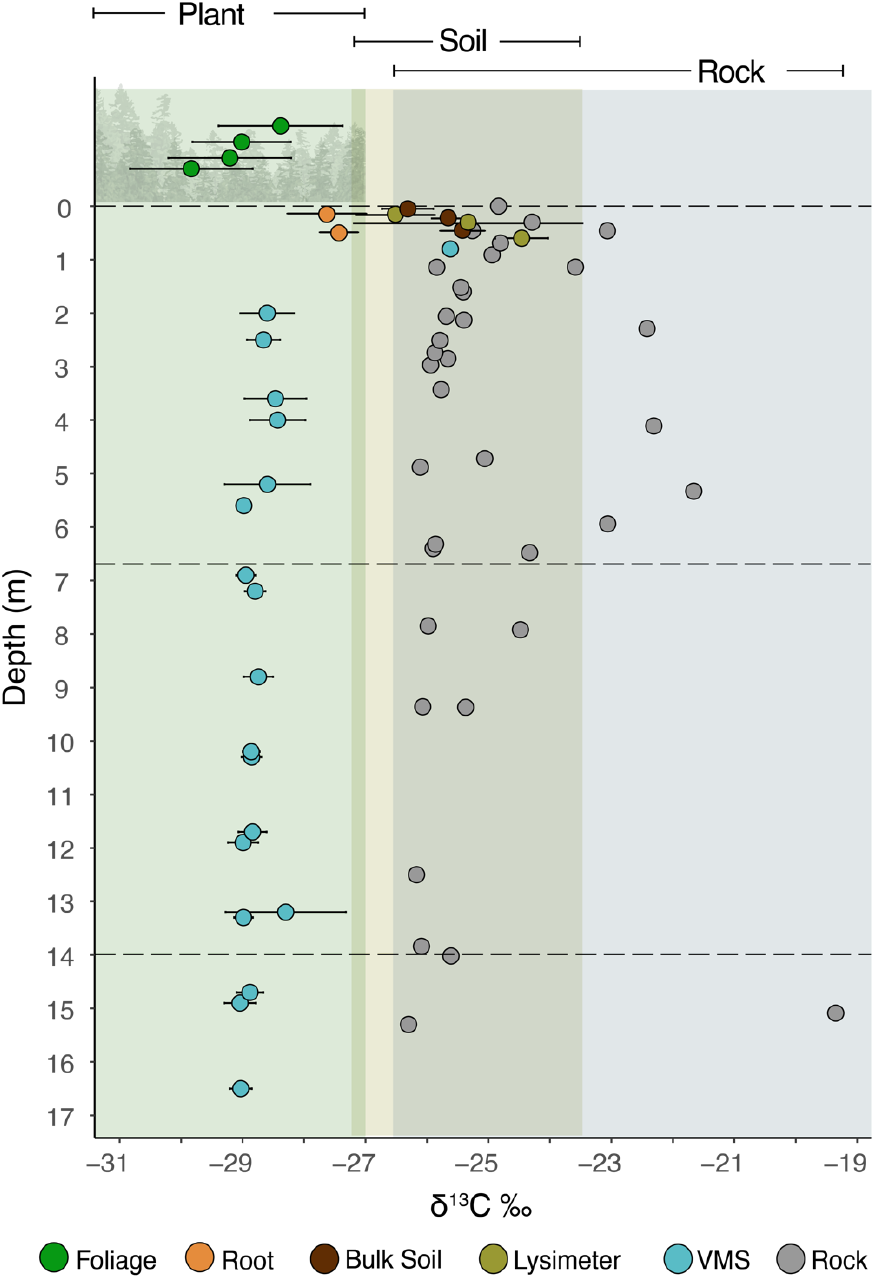
δ^13^C of DOC from VMS, soil lysimeters, foliage, root, builk soil and bulk rock. Foliage represents the four primary canopy trees (*Pseudotsuga menziesii, Notholithocarpus densiflorus, Arbutus menziesii, Quercus chrysolepis*), *Pseudotsuga menziesii* roots represent the root values. Where error bars exist, points represent an average across multiple samples. VMS samples are averaged over all δ^13^C-DOC sampling dates (see methods).

These findings indicate that the DOC source in the VMS is most consistent with minimally processed C_3_ plant material. Due to the strong link of DOC and TDN, this means that DON in the system is also of modern plant origin, and not from the weathering of the N-rich bedrock as has been observed in other systems (25, 28). Consistently low concentrations of NH_4_^+^ in the VMS further support the hypothesis that rock-derived N is unlikely to be a significant source of N to the WBVZ, as NH_4_^+^ is commonly the form in which N weathers from rock (34). However, organic N can also weather out of rock directly, but CO_2_ radiocarbon dating at the site found that respired carbon in the WBVZ was of modern plant origin and not petrogenic (35).

## Discussion

For the first time, we document significant amounts of biologically available N cycled within and below an active weathered bedrock rhizosphere year-round. We investigate the source and fate of TDN and its linked cycling with DOC to infer ecosystem-level drivers of N cycling within the WBVZ. We find that TDN concentrations are influenced by periods of high metabolic activity and inter-annual differences in precipitation patterns at the site, but that these patterns are also strongly influenced by depth in the WBVZ. Ultimately, we propose that plant activity within and below the rooting zone (0–7 m) drives linked TDN and DOC dynamics, shaping the source and the fate of dissolved organic matter in the WBVZ.

### Plant-derived organic matter shapes N cycling in the WBVZ

Within the WBVZ, TDN was primarily in the form of DON and strongly correlated with DOC, resulting in a consistent DOC:TDN ratio across all depths. Although dissolved organic matter in the WBVZ is very likely a complex mixture of compounds, this highly constrained DOC:TDN ratio of approximately 11 suggests several classes of organic molecules with similar C:N ratios may dominate, such as: microbial residues (23, 36), rhizodeposits and other small organic compounds produced via decomposition of plant and microbial biomass (37). Stable carbon isotope analysis of DOC further suggests dissolved organic matter in the WBVZ to be plant-derived (Fig. 1A, S3). SOM and bulk soil δ^13^C typically become more enriched with depth due to various mechanisms, namely, increased contributions of microbially derived carbon (33). Were leaching from soil (S1) the main mechanism of dissolved organic matter delivery to the WBVZ, we would expect δ^13^C-DOC in the VMS to be isotopically enriched by several permil compared to litter inputs, as is seen in both bulk soil and soil pore water DOC at the base of the soil profile (Fig. 6). Likewise, if shale-derived organic matter were the dominant source (S2), we would also expect δ^13^C-DOC in the VMS to be more enriched, falling within the range expected for rock—although isotopic fractionation during weathering of petrogenic organic matter remains poorly constrained (38). Taken together, the isotopically depleted δ^13^C-DOC in the WBVZ and previous radiocarbon dating at the site suggest modern, plant-derived compounds are the primary source of dissolved organic matter to the WBVZ.

### Rhizosphere activity and annual precipitation influence TDN concentration in the WBVZ

Plant-derived organic matter as the primary source of DOC and TDN in the WBVZ is consistent with an active rhizosphere extending meters into rock. We observe that TDN and NH_4_^+^ concentrations are lowest within the rooting zone—where we see a strong respiration signal—and accumulate with depth, likely resulting from higher rates of uptake in the shallow rooting zone and reduced rates of metabolic activity and uptake in progressively deeper horizons. We propose rhizosphere uptake drives the decrease in TDN concentrations between spring and summer in 2018, when leaching is likely low (Fig. 5) and forest-level metabolic activity is increasing (29). Uptake is also a likely mechanism of decrease in NH_4_^+^ between spring and summer months, as it is not only a highly bioavailable form of N, but also less susceptible to leaching as a cation in negatively charged soils. Decrease in TDN is very apparent in the rooting zone, however, we also observe a decrease in the deeper WBVZ as well, where we predict root abundance and activity to be very low. Such patterns may be the result of deep microbial activity, where microbes not strictly part of the rhizosphere utilize dissolved organic matter (39). But due to the convergent timing of plant metabolic activity and end of the wet season in the mediterranean climate of the ERCZO, separating the competing processes of uptake (F1) and leaching (F3) in the fate of TDN is challenging, especially between years as precipitation regimes change.

Timing and quantity of precipitation between 2018 and 2019 also had a strong influence on TDN concentration within the WBVZ. While we observed a systematic decrease in TDN from spring to summer months in 2018, we did not see the same in 2019. We suggest large quantities of rain received in winter 2019 led to spikes and ultimate loss through leaching of TDN from WBVZ prior to the onset of the growing season. Other studies have found that high rates of precipitation can result in large amounts of water entering and flushing through the vadose zone, increasing discharge at the stream and concomitant N losses (40, 41). We hypothesize that increased N residence times in the WBVZ during 2018 led to higher rates of N immobilization by plants and microbes, but that rapid loss of TDN in winter 2019 led to consistently low concentrations throughout the growing season.

Perhaps another indication of leaching as a mechanism of N loss is the low concentrations of NO_3_^-^ at the site. Unlike NH_4_^+^, NO_3_^-^ levels remained very low across all depths in the VMS throughout the sampling period. This is in keeping with most acidic, negatively charged temperate forest soils where NO_3_^-^ is readily lost via leaching. Another explanation could be denitrification. But given most of the vadose zone is unsaturated and oxic much of the year, denitrification is likely very low at the site, or restricted to anoxic microsites and deeper horizons where seasonally and chronically saturated bedrock may become anoxic, leading to hot spots and hot moments of N_2_O production under reducing conditions (42). However, as 95% of TDN is in the form of DON, on average, and the prevailing conditions are unfavorable, we predict denitrification not to be a major pathway of N loss at the site. Instead, we conclude plant and microbial use of inorganic and organic N residues, as well as leaching under high precipitation account for changes in TDN observed in the WBVZ.

### The Importance of the Deep N Cycle

Both source and fate of N in ecosystems remain challenging to study due to myriad competing processes, pools, and sinks (43). The quantity, composition, and seasonal dynamics of N in weathered bedrock reported here, suggest it plays a critical and overlooked role in nutrient ecology of terrestrial ecosystems, and represents an important pool of N not currently represented in ecosystem models. In ecosystems where soils are shallow or non-existent, the dynamic cycling of nutrients in weathered bedrock may be as important as those in soils. This challenges the long-standing paradigm that the majority of plant nutrient uptake occurs exclusively within soils. Our data clearly demonstrate that roots are biogeochemically active and play key functional roles in the weathered bedrock, extending important biogeochemical cycles to the base of the critical zone (12). To this end, we propose a revision to the standard ‘textbook’ version of the N cycle to include cycling in weathered bedrock where deep rhizospheres are known to be present.

It is critical that future research explores N cycling within weathered bedrock rhizospheres across diverse ecosystems. We predict the N cycle within the WBVZ is highly dependent on ecosystem characteristics, such as lithology, vegetation type, climate, depth to the groundwater table and subsurface water storage dynamics (44, 45). We put forward a conceptual model to describe pools and fluxes within the WBVZ that drive N cycling (Fig. 1A), noting that the magnitudes of these pools and fluxes will likely change when comparing weathered bedrock rhizospheres in distinct ecosystems. For example, the only other study documenting deep N in a WBVZ reports very different sources and fates of N. In their study, Wan et al. (28), found significant amounts of rock-derived N in the WBVZ primarily in the form of NO_3_^-^, and recorded major losses of N via denitrification (27). Importantly, their study site lacks a deep rhizosphere and the water table rises to the ground surface annually, making the vadose zone conditions quite different from those of the ERCZO. Future research is needed to understand variation in WBVZ N cycling and how it is shaped by ecosystem characteristics. Particularly, we propose cross-ecosystem study using deep monitoring systems like those deployed at the ERCZO and East River for monitoring of deep N dynamics in response to important ecosystem factors.

Our data support the assertion that the fractured bedrock zone is a reservoir, not only for water, but essential nutrients that are cycled by deeply rooted vegetation. In the future, as N resources become even more limiting with seasonal shifts in water availability under more frequent and severe droughts (52), rising temperatures and vegetation changes, this deep resource reservoir is likely to become increasingly important for the growth and physiology of deeply rooted plants. Our research highlights the need for future critical zone research to expand N cycling measurements and modeling to include weathered bedrock horizons. Only then can we begin to understand the extent to which forests and other deeply rooted ecosystems utilize deep nutrient stores and uncover the roles of weathered bedrock rhizospheres across diverse environments.

## Methods

### Field Site

Our study site (Rivendell, 39.729 N, 123.644 W) is an intensively monitored (https://dendra.science/orgs/erczo), steep (average 32°), north-facing hillslope within the ERCZO (Figure 1 panel A) hosted at the UC Angelo Coast Range Reserve in northern California. At the site, soils mantle a deeply weathered argilicious bedrock mapped as marine turbidites of the Coastal Belt of the Franciscan Complex (46). Rivendell is home to a Vadose Zone Monitoring System (VMS; Figure 1, panel B), which enables discrete sampling of water, gasses, temperature, and relative rock moisture content at approximately 1.5-meter intervals down to approximately 16 meters depth (47). This is the only VMS that has been installed exclusively for long-term environmental monitoring of an old growth forested ecosystem and offers a unique look into the deeply weathered bedrock rhizosphere underlying the Rivendell hillslope.

The vegetation at the site is characterized as a mixed needleleaf-broadleaf evergreen forest. The dominant species include Douglas fir (*Pseudotsuga menziesii*), live oak (*Quercus chrysolepis)*, tan oak (*Notholithocarpus densiflorus*) and madrone (*Arbutus menziesii*). Woody vegetation rooting profiles are estimated to extend to approximately 7 meters depth (29) and peak transpiration rates typically occur in early July (30).

### Vadose zone water and gas sampling

The VMS consists of two 55° inclined boreholes installed along contour on the N-facing Rivendell hillslope (Fig. 1B). Both boreholes extend down to the water table at approximately 16 meters depth below the ground surface. These boreholes are lined with flexible sleeves instrumented with sensors and sampling ports designed to capture either freely draining (Sleeve B) or matrix held water (Sleeve A) using passive or tension-induced sampling methods (Fig. 1C), technical methodology of the VMS is outlined in detail elsewhere (47–49). Gas ports are embedded within the feely draining moisture sampling ports along the length of sleeve B to also allow for sampling of vadose zone gasses. Ports 1–4 (< 6.9 m) are located within the chronically unsaturated vadose zone, while ports 5–9 (7-14.9 m) are seasonally saturated and the deepest port (port 10, > 15 m) is chronically saturated for both sleeves (Figure 1C).

### VMS sampling

We sampled the VMS from January of 2018 to January of 2020 for Total Dissolved N (TDN) and Dissolved Organic Carbon (DOC) at approximately bi-monthly intervals. Inorganic N (NO_3_^-^ and NH_4_^+^) sampling commenced in July of 2018 and continued to March 2020 (Figure S1). Pore water samples in the VMS were collected by applying a light vacuum (−0.7 bar) to all ports and allowing water to be drawn to ports over approximately 2 weeks in between sampling periods, as described in previous studies at the site (29). All water samples were passed through a 0.2-micron filter in the field immediately upon sampling. Water samples for NH_4_^+^ and NO_3_^-^ were stored in acid-washed HDPE bottles at -20ºC prior to analysis by the diffusion-conductivity method (UC Davis Analytical Laboratory). Water samples for TDN and DOC analysis were filtered into cleaned and baked glass vials with Teflon caps and stored at 4ºC prior to analysis. TDN and DOC concentrations (± 0.001 mg/L) were quantified by either a Teledyne Tekmar Apollo 9,000 Carbon Analyzer (UT Jackson School of Geosciences Aqueous Geochemistry Lab) or an Injection Flow Analyzer (UC Davis Analytical Laboratory).

Additionally, DOC was analyzed for δ^13^C from a subset of sampling dates in 2021 in the VMS (March 6, May 16, June 2, August 24, July 25) and soil pore water samplers (SK20 pore water sampler, Meter Group, Inc) when soils were wet enough for samples to be pulled by light vacuum (January 1, February 2, March 6). δ^13^C-DOC analyses were performed at the Stable Isotope Facility at University California, Davis, using an O.I. Analytical Model 1030 TOC Analyzer (Xylem Analytics, College Station, TX), interfaced with a PDZ Europa 20-20 isotope ratio mass spectrometer (Sercon Ltd., Cheshire, UK) outfitted with a GD-100 Gas Trap Interface (Graden Instruments). Filtered DOC samples were stored at -20ºC prior to δ^13^C analysis.

We calculate dissolved organic N (DON) from TDN and inorganic N measurements ([DON] = [TDN] - ([NH_4_^+^] + [NO_3_^-^]). Due to the oxic and slightly acidic (pH ∼ 6) nature of the ERCZO vadose zone, we estimate total inorganic N to be NH_4_^+^ and NO_3_^-^, as other inorganic N forms (e.g. NO_2_^-^ and NH_3_) are often below detection (50, 51). Additionally, CO_2_ and O_2_ concentrations were measured (Quantek gas analyzer) from the VMS gas ports and one soil port (installed ∼30cm below soil surface) within 24 hours of water sampling for all sampling campaigns and always during daylight hours.

### Solids Sampling

Foliage was sampled in summers 2018 and 2019 from canopies of four species of dominant tree species at the Eel River CZO: *Arbutus menziesii, Pseudotsuga menziesii, Quercus chrysolepis* and *Notholithocarpus densiflorus*. All foliage samples were dried at 60ºC for elemental and stable isotope analysis (at the Center for Stable Isotope Biogeochemistry, UC Berkeley). Soil cores were taken until the auger could no longer penetrate the subsurface any further (max depth ∼30 cm–1 m deep) across the hillside surrounding the VMS. Gravimetric soil moisture content as well as elemental and stable isotope analyses were quantified for these soil samples. Rock samples were collected during the installation of the VMS via deep bore holes. Samples were dried, pulverized and analyzed for total elemental composition and stable isotope analysis at the UC Davis Stable Isotope Facility (EA-IRMS).

### Statistical Analyses

We used linear mixed models (*lme4* package) in R (v4.2.2) to explore the effects of season, depth, year, annual cumulative precipitation (fixed effects) and Port ID (random effect) on TDN and NH_4_^+^ concentrations in the VMS (52, 53). NO_3_^-^ was not included in these analyses as it is frequently below detection. We conducted model selection using Akaike’s information criterion (AIC) to estimate model goodness-of-fit. Models for each response variable that had the lowest AIC were selected to perform further statistical analyses, including Type II ANOVA in the *lmerTest* package to test significance of fixed effects on TDN and NH_4_^+^ (54). Pairwise comparisons were conducted using the *emmeans* package in R for post hoc Tukey comparison of seasonal differences in TDN within water year (55) and the *r*.*squaredGLMM* command from MuMIn package was used to calculate marginal and conditional R^2^ for all models (56). Violations of homogeneity of variance and linearity assumptions were tested for all final models using the *check_model* command from the *performance* package in R (57).

For simplicity, we refer to the time periods as distinct seasons from here on out: spring (Feb–Apr), summer (May–July) and winter (Nov–Jan). We define these cutoffs based on observations of vegetation phenology (i.e. timing of bud break and peak transpiration rates) as well as patterns in seasonal precipitation at the site. Ultimately, spring months represent the early growing season at the site and summer months represent mid-growing season at the site where the WBVZ is still largely wet enough for sampling of pore water to occur. We include winter values (Nov–Jan) in our full model of TDN to capture all seasons where the VMS is wet enough for sampling, but exclude Fall months as the shallowest portions of the VMS are too dry to sample water from. We also refer to the water year (e.g. Oct 2017–Oct 2018 for 2018 water year) instead of year of sampling alone, as it more accurately captures annual variation in precipitation and site dynamics. We focus on water years 2018 and 2019 for linear mixed models.

## Acknowledgments

We would like to acknowledge all of our collaborators at the Eel River Critical Zone Observatory for all of their support over the years. We would particularly like to thank Bill Dietrich and Jenny Druhan for their guidance and support throughout the course of this project. We would also like to thank our field technicians Hunter Jamison, Gunnar Rieth, Will Speiser, Marshall Wolf and Samantha Cargill for their hard work sampling the VMS, as well as undergraduate field assistants that helped over the years. We thank Collin Bode for development and maintenance of site metadata and Wendy Baxter and Anthony Ambrose for their expert help in accessing tree canopies for sampling at the site. We also thank Scott Mitchell, Benjamin Houlton and Randy Dahlgren for providing rock carbon isotope data. We thank members of the Dawson Lab for feedback on the research presented here and early drafts of the manuscript. Thank you to Whendee Silver and David Ackerly for their extensive feedback on this manuscript and mentorship for many years. Thank you to Stefania Mambelli and Wenbo Yang for running samples for stable isotope analysis at the Center For Stable Isotope Biogeochemistry and UC, Berkeley. Thank you to the UC Davis Analytical Lab for analytical support. We would also like to thank our colleagues, Benjamin R Karin and Jesus Martinez-Gomez for feedback on manuscript drafts. And lastly, to all the incredible field and lab assistants that made this work possible: Saumitra Kelkar, Hannah Johansson, Alexandra Carey and Olivia Parra. Thank you.

## References

1. J.-L. Maeght, B. Rewald, A. Pierret, How to study deep roots—and why it matters. Front. Plant Sci. 4 (2013).

2. R. B. Jackson, et al., A global analysis of root distributions for terrestrial biomes. Oecologia 108, 389–411 (1996).

3. P. Poot, H. Lambers, Shallow-soil endemics: adaptive advantages and constraints of a specialized root-system morphology. New Phytol. 178, 371–381 (2008).

4. H. J. Schenk, Soil depth, plant rooting strategies and species’ niches. New Phytol. 178, 223–225 (2008).

5. S. Schwinning, The ecohydrology of roots in rocks. Ecohydrology 3, 238–245 (2010).

6. M. A. Zwieniecki, M. Newton, Seasonal pattern of water depletion from soil–rock profiles in a Mediterranean climate in southwestern Oregon. Can. J. For. Res. 26, 1346–1352 (1996).

7. D. M. Rempe, W. E. Dietrich, Direct observations of rock moisture, a hidden component of the hydrologic cycle. Proc. Natl. Acad. Sci. 115, 2664–2669 (2018).

8. E. L. McCormick, et al., Widespread woody plant use of water stored in bedrock. Nature 597, 225–229 (2021).

9. T. E. Dawson, W. J. Hahm, K. Crutchfield-Peters, Digging deeper: what the critical zone perspective adds to the study of plant ecophysiology. New Phytol. 226, 666–671 (2020).

10. B. C. McLaughlin, et al., Hydrologic refugia, plants, and climate change. Glob. Change Biol. 23, 2941–2961 (2017).

11. P. Hinsinger, C. Plassard, B. Jaillard, Rhizosphere: A new frontier for soil biogeochemistry. J. Geochem. Explor. 88, 210–213 (2006).

12. D. de B. Richter, S. A. Billings, ‘One physical system’: Tansley’s ecosystem as Earth’s critical zone. New Phytol. 206, 900–912 (2015).

13. D. S. LeBauer, K. K. Treseder, Nitrogen Limitation of Net Primary Productivity in Terrestrial Ecosystems Is Globally Distributed. Ecology 89, 371–379 (2008).

14. T. Näsholm, K. Kielland, U. Ganeteg, Uptake of organic nitrogen by plants. New Phytol. 182, 31–48 (2009).

15. T. Kraiser, D. E. Gras, A. G. Gutierrez, B. Gonzalez, R. A. Gutierrez, A holistic view of nitrogen acquisition in plants. J. Exp. Bot. 62, 1455–1466 (2011).

16. H. Marschner, Marschner’s Mineral Nutrition of Higher Plants (Academic Press, 2011).

17. D. Moreau, R. D. Bardgett, R. D. Finlay, D. L. Jones, L. Philippot, A plant perspective on nitrogen cycling in the rhizosphere. Funct. Ecol. 33, 540–552 (2019).

18. A. C. Finzi, et al., Rhizosphere processes are quantitatively important components of terrestrial carbon and nutrient cycles. Glob. Change Biol. 21, 2082–2094 (2015).

19. L. Henneron, P. Kardol, D. A. Wardle, C. Cros, S. Fontaine, Rhizosphere control of soil nitrogen cycling: a key component of plant economic strategies. New Phytol. 228, 1269–1282 (2020).

20. M. M. M. Kuypers, H. K. Marchant, B. Kartal, The microbial nitrogen-cycling network. Nat. Rev. Microbiol. 16, 263–276 (2018).

21. A. Jilling, M. Keiluweit, J. L. M. Gutknecht, A. S. Grandy, Priming mechanisms providing plants and microbes access to mineral-associated organic matter. Soil Biol. Biochem. 158, 108265 (2021).

22. L. Philippot, S. Hallin, G. Börjesson, E. M. Baggs, Biochemical cycling in the rhizosphere having an impact on global change. Plant Soil 321, 61–81 (2009).

23. C. C. Cleveland, D. Liptzin, C:N:P stoichiometry in soil: is there a “Redfield ratio” for the microbial biomass? Biogeochemistry 85, 235–252 (2007).

24. R. A. Dahlgren, Soil acidification and nitrogen saturation from weathering of ammonium-bearing rock. Nature 368, 838–841 (1994).

25. S. L. Morford, B. Z. Houlton, R. A. Dahlgren, Direct quantification of long-term rock nitrogen inputs to temperate forest ecosystems. Ecology 97, 54–64 (2016).

26. B. Z. Houlton, S. L. Morford, R. A. Dahlgren, Convergent evidence for widespread rock nitrogen sources in Earth’s surface environment. Science 360, 58–62 (2018).

27. E. A. Hasenmueller, et al., Weathering of rock to regolith: The activity of deep roots in bedrock fractures. Geoderma 300, 11–31 (2017).

28. J. Wan, et al., Bedrock weathering contributes to subsurface reactive nitrogen and nitrous oxide emissions. Nat. Geosci. 14, 217–224 (2021).

29. A. K. Tune, J. L. Druhan, J. Wang, P. C. Bennett, D. M. Rempe, Carbon Dioxide Production in Bedrock Beneath Soils Substantially Contributes to Forest Carbon Cycling. J. Geophys. Res. Biogeosciences 125, e2020JG005795 (2020).

30. P. Link, et al., Species differences in the seasonality of evergreen tree transpiration in a Mediterranean climate: Analysis of multiyear, half-hourly sap flow observations. Water Resour. Res. 50, 1869–1894 (2014).

31. M. Christ, Y. Zhang, G. E. Likens, C. T. Driscoll, Nitrogen Retention Capacity of a Northern Hardwood Forest Soil Under Ammonium Sulfate Additions. Ecol. Appl. 5, 802–812 (1995).

32. Z. Yu, et al., Contribution of amino compounds to dissolved organic nitrogen in forest soils. Biogeochemistry 61, 173–198 (2002).

33. B. Boström, D. Comstedt, A. Ekblad, Isotope fractionation and 13C enrichment in soil profiles during the decomposition of soil organic matter. Oecologia 153, 89–98 (2007).

34. G. E. Bebout, K. E. Lazzeri, C. A. Geiger, Pathways for nitrogen cycling in Earth’s crust and upper mantle: A review and new results for microporous beryl and cordierite. Am. Mineral. 101, 7–24 (2016).

35. A. K. Tune, J. L. Druhan, C. R. Lawrence, D. M. Rempe, Deep root activity overprints weathering of petrogenic organic carbon in shale. Earth Planet. Sci. Lett. 607, 118048 (2023).

36. L. Sun, M. Ataka, Y. Kominami, K. Yoshimura, K. Kitayama, A constant microbial C/N ratio mediates the microbial nitrogen mineralization induced by root exudation among four co-existing canopy species. Rhizosphere 17, 100317 (2021).

37. P. Bengtson, J. Barker, S. J. Grayston, Evidence of a strong coupling between root exudation, C and N availability, and stimulated SOM decomposition caused by rhizosphere priming effects. Ecol. Evol. 2, 1843–1852 (2012).

38. T. Roylands, et al., Capturing the short-term variability of carbon dioxide emissions from sedimentary rock weathering in a remote mountainous catchment, New Zealand. Chem. Geol. 608, 121024 (2022).

39. P. Priyadharsini, et al., “Mycorrhizosphere: The Extended Rhizosphere and Its Significance” in Plant-Microbe Interaction: An Approach to Sustainable Agriculture, D. K. Choudhary, A. Varma, N. Tuteja, Eds. (Springer Singapore, 2016), pp. 97–124.

40. F. Hagedorn, J. B. Bucher, P. Schleppi, Contrasting dynamics of dissolved inorganic and organic nitrogen in soil and surface waters of forested catchments with Gleysols. Geoderma 100, 173–192 (2001).

41. R. B. Alexander, et al., Dynamic modeling of nitrogen losses in river networks unravels the coupled effects of hydrological and biogeochemical processes. Biogeochemistry 93, 91–116 (2009).

42. M. E. McClain, et al., Biogeochemical Hot Spots and Hot Moments at the Interface of Terrestrial and Aquatic Ecosystems. Ecosystems 6, 301–312 (2003).

43. S. Niu, et al., Global patterns and substrate-based mechanisms of the terrestrial nitrogen cycle. Ecol. Lett. 19, 697–709 (2016).

44. H. Jenny, Factors of soil formation: a system of quantitative pedology (Dover, 1941).

45. H. Jenny, Role of the Plant Factor in the Pedogenic Functions. Ecology 39, 5–16 (1958).

46. R. J. McLaughlin, et al., Geology of the Cape Mendocino, Eureka, Garberville, and Southwestern part of the Hayfork 30 x 60 Minute Quadrangles and Adjacent Offshore Area, Northern California, Miscellaneous Field Studies Map MF‐2336. USGS, Washington, DC (2000).

47. J. L. Druhan, N. Fernandez, J. Wang, W. E. Dietrich, D. Rempe, Seasonal shifts in the solute ion ratios of vadose zone rock moisture from the Eel River Critical Zone Observatory. Acta Geochim. 36, 385–388 (2017).

48. Y. Rimon, O. Dahan, R. Nativ, S. Geyer, Water percolation through the deep vadose zone and groundwater recharge: Preliminary results based on a new vadose zone monitoring system. Water Resour. Res. 43 (2007).

49. O. Dahan, et al., In Situ Monitoring of Water Percolation and Solute Transport Using a Vadose Zone Monitoring System Vadose Zone J. 8, 916–925 (2009).

50. O. Cleemput, A. H. Samater, Nitrite in soils: accumulation and role in the formation of gaseous N compounds. Fertil. Res. 45, 81–89 (1995).

51. A. O. Langford, F. C. Fehsenfeld, J. Zachariassen, D. S. Schimel, Gaseous ammonia fluxes and background concentrations in terrestrial ecosystems of the United States. Glob. Biogeochem. Cycles 6, 459–483 (1992).

52. D. Bates, M. Mächler, B. Bolker, S. Walker, Fitting Linear Mixed-Effects Models Using lme4. J. Stat. Softw. 67, 1–48 (2015).

53. R Core Team, R: A language and environment for statistical computing. R Foundation for Statistical Computing, Vienna, Austria. ## URL https://www.R-project.org/. (2022).

54. A. Kuznetsova, P. B. Brockhoff, R. H. B. Christensen, lmerTest Package: Tests in Linear Mixed Effects Models. J. Stat. Softw. 82 (2017).

55. R. V. Lenth, et al., emmeans: Estimated Marginal Means, aka Least-Squares Means (2023) (April 19, 2023).

56. K. Barton, Package ‘mumin.’ Version 1, 439 (2015).

57. D. Lüdecke, M. S. Ben-Shachar, I. Patil, P. Waggoner, D. Makowski, performance: An R package for assessment, comparison and testing of statistical models. J. Open Source Softw. 6 (2021).

