## Supplemental Materials for "Deep rhizospheres extend the nitrogen cycle meters below the base of soil into weathered bedrock"

**Sample Collection Note**: TDN data were collected approximately on a bi-weekly basis from January 2018 until January 2020. DOC was also sampled using this same schedule, however analysis stopped in May of 2019 due to instrument failure. Ammonium (NH4+) and nitrate (NO3-) analyses began in May of 2018 are were always run together until March 2020.

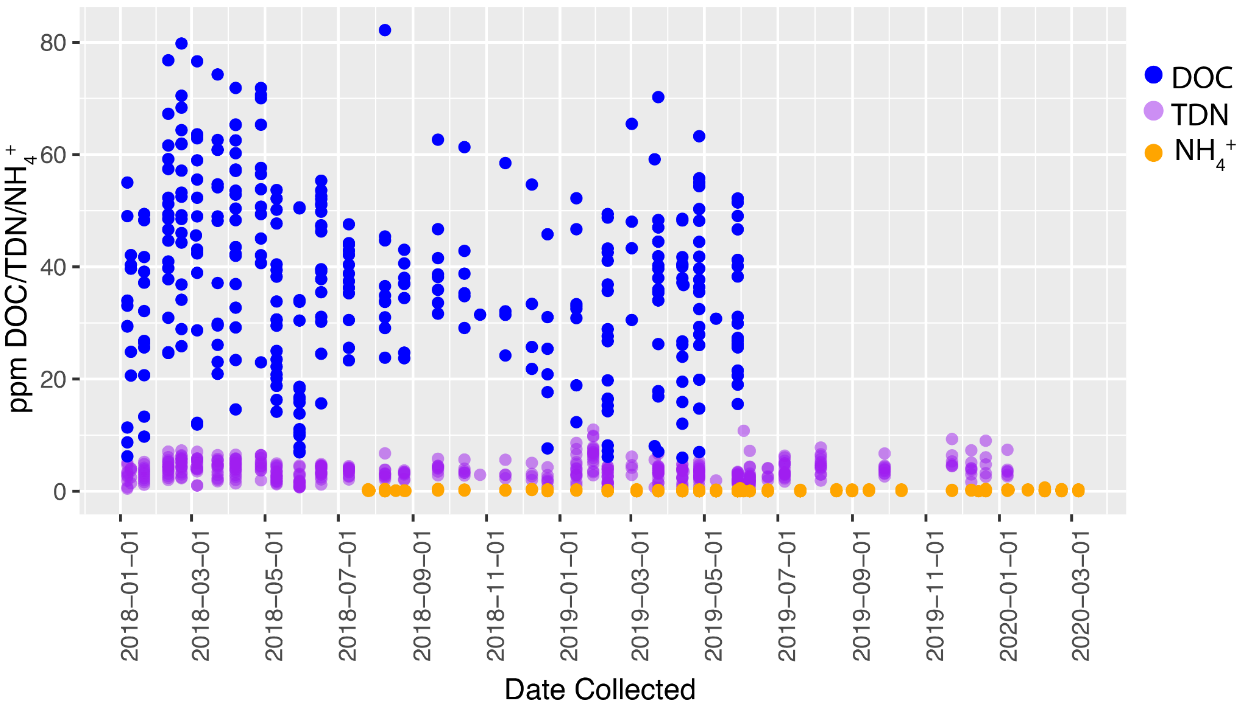


**Figure S1**. Sample Collection Date ranges, overlap and gaps. Dissolved organic carbon (DOC; blue), total dissolved nitrogen (TDN; purple), and ammonium (NH4+; orange—representing both NH4+ and NO3- collection dates).

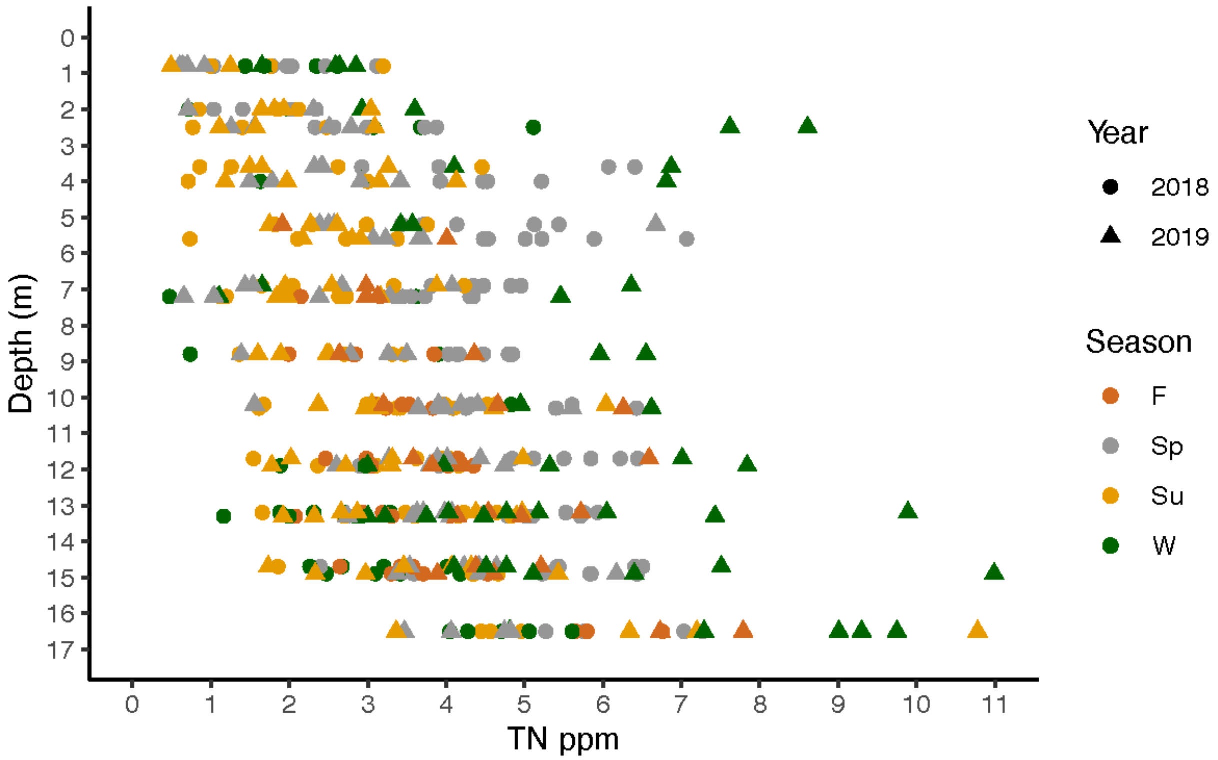


**Figure S2.** Total Dissolved Nitrogen for all sampling dates in the Vadose Zone Monitoring system (VMS) across the entire VMS depth profile. Colors indicate season, shape indicates year.

**Table S1**. Summary of Type II ANOVA results for Full model of Total Dissolved Nitrogen *lmer* model: (TDN~Depth + Year*Season + (1|PortID)

| Parameter | Sum Sq | Mean Sq | Num DF | DF | F value | Pr(>F) |
|---|---|---|---|---|---|---|
| Depth | 66.2 | 66.2 | 1 | 16.3 | 39.7 | **<0.0001** |
| Season | 99.7 | 49.9 | 2 | 378.1 | 29.9 | **<0.0001** |
| Water year | 11.4 | 11.4 | 1 | 374.1 | 6.8 | 0.0093 |
| Season:Water year | 95.4 | 47.7 | 2 | 374.9 | 28.6 | **<0.0001** |

Degrees-of-freedom method: Kenward-Roger

**Table S2.** Summary of *emmeans* for 2018 and 2019 water years by season

| Water Year | Season | emmean | Standard Error | DF | Lower CL | Upper CL |
|---|---|---|---|---|---|---|
| 2018 | | | | | | |
| | Spring | 4.56 | 0.16 | 62.23 | 4.23 | 4.89 |
| | Summer | 2.96 | 0.18 | 91.09 | 2.60 | 3.32 |
| | Winter | 3.59 | 0.22 | 142.14 | 3.15 | 4.02 |
| 2019 | | | | | | |
| | Spring | 3.18 | 0.18 | 83.68 | 2.83 | 3.53 |
| | Summer | 2.93 | 0.19 | 105.68 | 2.55 | 3.31 |
| | Winter | 4.69 | 0.22 | 152.68 | 4.26 | 5.12 |

Degrees-of-freedom method: Kenward-Roger
Confidence level used: 0.95

**Table S3**. Summary of estimated marginal means from pairwise comparison of TDN by season for 2018 and 2019 water years using *emmeans* package

| Water year | Contrast | Estimate | Standard Error | DF | t.ratio | p value |
|---|---|---|---|---|---|---|
| 2018 | | | | | | |
| | Sp - Su | 1.595 | 0.20 | 372.94 | 7.83 | **<0.0001** |
| | Sp - W | 0.972 | 0.24 | 381.53 | 4.12 | **0.0001** |
| | Su - W | -0.624 | 0.25 | 383.32 | -2.49 | **0.0350** |
| 2019 | | | | | | |
| | Sp - Su | 0.251 | 0.22 | 371.92 | 1.13 | 0.4956 |
| | Sp - W | -1.515 | 0.25 | 375.83 | -6.16 | **<0.0001** |
| | Su - W | -1.766 | 0.26 | 375.29 | -6.92 | **<0.0001** |

Sp = Spring, Su = summer, W = Winter

**Table S4**. Summary of Type II ANOVA results for Full model of NH4+ concentrations
*lmer* model: (TDN~Depth + Season + (1|PortID)

| Parameter | Sum Sq | Mean Sq | NumDF | Den DF | F value | Pr(>F) |
|---|---|---|---|---|---|---|
| Season | 0.15 | 0.15 | 1 | 213.88 | 28.4 | **< 0.0001** |
| Depth | 0.18 | 0.18 | 1 | 17.54 | 35.2 | **< 0.0001** |

Degrees-of-freedom method: Kenward-Roger

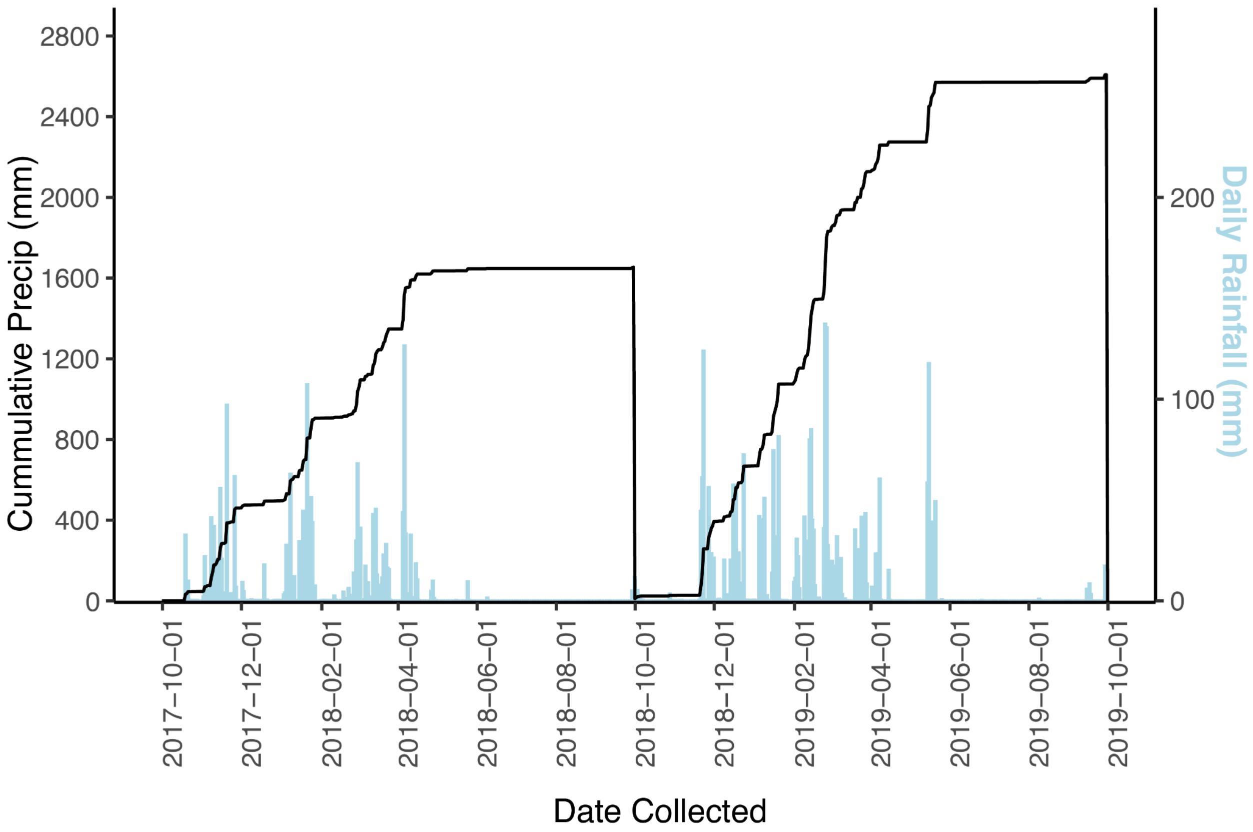


**Figure S3**. Cumulative precipitation (mm, black line, left y-axis) and daily rainfall (mm, blue bars, right y-axis) for the duration of the sampling period.

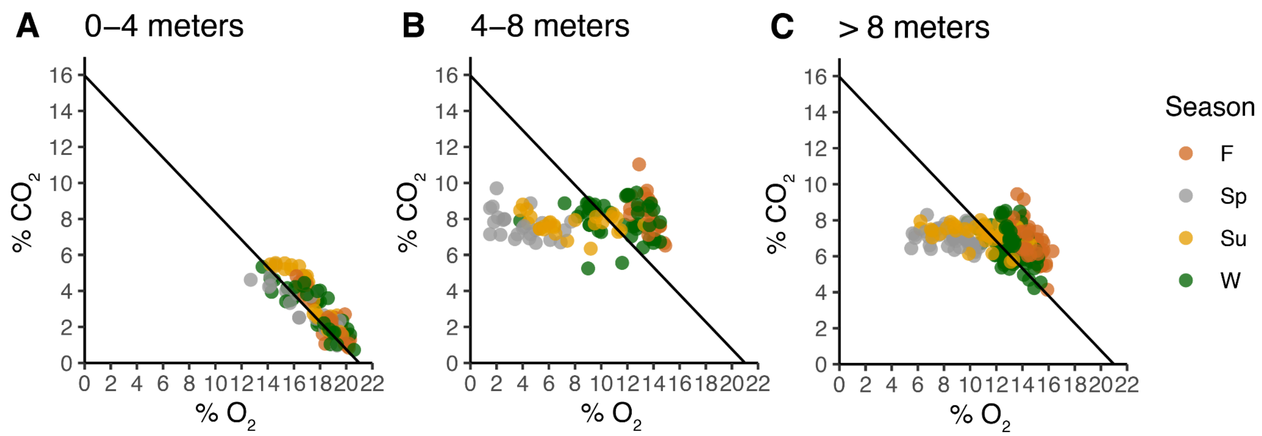


**Figure S4**. $CO_2$ to $O_2$ concentrations within the first 4 m, 4-8 m, > 8 m from the soil surface during samples bi-weekly from October 2017-December 2019. The black solid line represents a reference respiratory quotient for aerobic respiration a 1:1 stoichiometric relationship between $O_2$ consumption and $CO_2$ production (RQ = 1) corrected for the differences in diffusivity coefficients between $O_2$ and $CO_2$ so that the slope is -0.76 with an x intercept = 21 (atmospheric $O_2$ concentration). These data include a subset of $CO_2$ and $O_2$ values originally reported in Tune et al. (1)(Jan 2018-Apr 2019) but include an additional 8 months from 2019 (May 2019- Dec 2019). While 0-4 meters falls consistently along the line representing respiration, below 4 m depth other processes ($CO_2$ dissolution and possibly mineral oxidation) compete with respiration in driving the $CO_2$ to $O_2$ relationship.

### References


1. A. K. Tune, J. L. Druhan, J. Wang, P. C. Bennett, D. M. Rempe, Carbon Dioxide Production in Bedrock Beneath Soils Substantially Contributes to Forest Carbon Cycling. *J. Geophys. Res. Biogeosciences* **125**, e2020JG005795 (2020).
